# Assembly archetypes in ecological communities

**DOI:** 10.1101/2023.06.20.545780

**Authors:** Hugo Flores-Arguedas, Omar Antolin-Camarena, Serguei Saavedra, Marco Tulio Angulo

## Abstract

An instrumental discovery in comparative and developmental biology is the existence of assembly archetypes that synthesize the vast diversity of organisms’ body plans—from legs and wings to human arms—into simple, interpretable, and general design principles. Here we combine a novel mathematical formalism based on Category Theory with experimental data to show that similar “assembly archetypes” exist at the larger organization scale of ecological communities when assembling a species pool across diverse environmental contexts, particularly when species interactions are highly structured. We applied our formalism to clinical data discovering two assembly archetypes that differentiate between healthy and unhealthy human gut microbiota. The concept of assembly archetypes and the methods to synthesize them can pave the way to discovering the general assembly principles of the ecological communities we observe in nature.

## Introduction

The ultimate goal of biology is to synthesize the general design principles that underlie the diversity of life on Earth [1]. A very insightful example of such synthesis in comparative and developmental biology is the discovery of assembly archetypes that synthesize the diversity of organisms’ body plans into simple, interpretable, and general design principles [2–5]. To illustrate such assembly archetypes, consider the wings that birds use to fly, the legs that mice use to run, and the arms that we humans use to create works of art (Fig. 1a). These limb *types* have a distinct number of bones, dimensions, and functions. Yet, underlying these differences, their body plans share the same *assembly archetype* of tetrapods [4, 5]: one (proximal stylopod) bone, followed by two (middle zeugopod) bones, ending with distal (autopod) bones. An assembly archetype can also be the substrate to build new, more complex assembly rules and assembly archetypes [6]. In our example, requiring that distal bones are *digits* generates a more complex assembly archetype that synthesizes the limbs of mice and humans. These assembly archetypes of different complexity organize diverse limb types into a hierarchy that explains how they are related (Fig. 1b). Namely, a collection of limb types share an assembly archetype (i.e., a common design principle) if they have a common ancestor. For example, the green, yellow and purple types in Fig. 1b share an assembly archetype, but not the red type.

**Figure 1:**
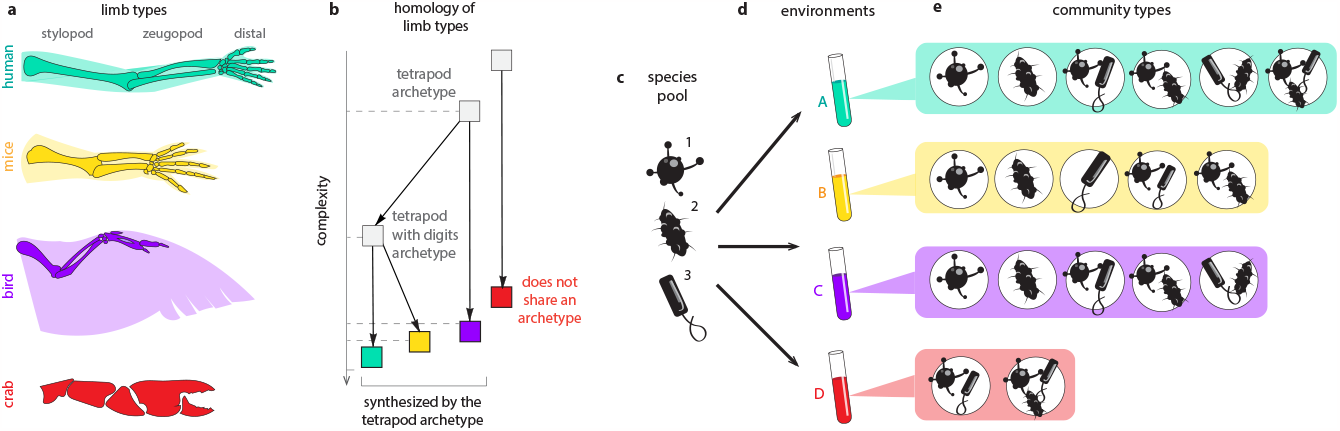
Assembly archetypes in ecological communities across environmental contexts. We introduce our approach using an analogy to how comparative and developmental biologists synthesize assembly archetypes in organisms’ body plans. **a**. Cartoons of the skeletons of four limb types. Despite differences in their size, the number of bones, and use, direct inspection shows that the first three limb types share the assembly archetype of tetrapods [4]: one (stylopod) bone, followed by two (zeugopod) bones, ending in distal bones. This archetype provides a simple, general, and interpretable assembly principle for different limb types. One can obtain the more complex assembly archetype “tetrapods with digits” starting from the “tetrapods” archetype and adding the constraint “distal bones are digits”. Importantly, note that limb types have endowed a structure allowing us to directly perceive their assembly archetypes by inspection using our human senses. **b**. The two assembly archetypes of the panel a (gray) of different complexity organize the limb types (colors) into a hierarchy. A collection of limb types have a common assembly principle if they share an assembly archetype, synthesized by their closest common ancestor in this hierarchy. In particular, limb types are more similar if the assembly archetype they share is closer in this hierarchy. Therefore, human and mouse limbs are more similar compared to bird limbs. Limb types cannot be synthesized into an assembly archetype when they do not share a common design principle (i.e., they do not share an ancestor in the hierarchy). This situation happens when we try to synthesize an assembly archetype for humans, mice, birds, and crabs in our example. **c**. Cartoon for a species pool Σ = {1, 2, 3} of *S* = 3 species. **d**. The species pool can be assembled in four hypothetical environmental contexts Θ = {*A, B, C, D*}. **e**. The coexistence of a species composition depends on the environmental context in which we assemble it. This fact generates different community types (i.e., different coexisting species compositions) across different environmental conditions. The assembly rule of an environment characterizes which community types we can observe in that environment. That is, just as limbs adopt different types in different organisms, the same species pool adopts different community types when assembled in different environmental contexts. However, unlike limb types, the community types come with no structure organizing them, making it virtually impossible to synthesize their assembly archetypes. For example, by inspecting the four community types in this panel, it is difficult for human senses to grasp if they share an assembly archetype or not.

In ecology, however, it is still unclear if general design principles underlie the assembly of ecological communities across the diverse environmental contexts found in nature [7–15]. Indeed, because species coexistence depends on many environmental factors such as temperature [16] or available nutrients [17], the same species pool can exhibit different species compositions when assembled under different environmental contexts. That is, just like limbs “adopt” different types of form in different organisms, a species pool, when assembled under different environmental contexts, can “adopt” different *community types*. Here, the community type *H*_*θ*_ in the environmental context *θ* is the set of all species compositions that we can assemble by combining species from the pool in that environment. The community type describes all species compositions that coexist and hence we can observe in a given environmental context (see Methods for a precise definition). Figure 1c-e illustrates this concept using a hypothetical pool of *S* = 3 species that can be assembled under four different environmental contexts Θ = {*A, B, C, D*}. For instance, the community adopts the type *H*_*B*_ = {{1}, {2}, {1, 3}, {2, 3}} under environment *B* and the type *H*_*D*_ = {{1, 3}, {1, 2, 3}} under environment *D*. The composition {1, 3} belongs to both community types, so species 1 and 3 can coexist when assembled in both environmental contexts. That the composition {1} belongs only to *H*_*B*_ means that species 1 can survive alone in the environmental context *B* but not in *D*.

To obtain a first-hand experience of the difficulties of identifying general design principles in the assembly of ecological communities, consider the following question: do the community types in Fig. 1e share an “assembly archetype” analogous to the tetrapod assembly archetype that the limb types in Fig. 1a-b share? This question is not asking if the community types have something in common regarding their species compositions, such as if one species composition appears in all of them. That would be analogous to asking if various limb types have a common bone. Instead, the question asks for an assembly archetype characterizing a fundamental similarity in the rules used to assemble all community types. If community types do exhibit assembly archetypes, their existence can pave the way to synthesizing the general assembly principles of ecological communities across diverse environmental conditions, which have remained elusive and controversial since Diamond’s seminal work more than 40 years ago [7, 10, 18, 19].

Answering the above question using our human senses alone is difficult because, unlike limb types that have additional topological structure telling which bones are adjacent, community types are plain collections of numbers lacking any structure. Without this structure, it is virtually impossible to determine whether or not two different community types are the product of similar assembly rules. Consequently, it becomes impossible to determine if a given collection of community types shares an assembly archetype. This difficulty is not limited to Figure 1. Instead, it is a fundamental challenge to discover general design principles in ecological communities assembled across diverse environmental contexts [20]. Namely, classic [21] and more recent [22] work has shown that communities can exhibit well-defined assembly rules and specific community types when assembled under similar environmental contexts. However, these rules do not translate well under more diverse environmental contexts [14, 19, 21, 23], making it unclear if and in which sense such rules could be an assembly archetype.

Here we address the above challenges by introducing a mathematical formalism based on Category Theory [24] that endows the collection of community types with the necessary structure to compare their assembly rules. This formalism provides an abstract “synthesis operation” that we can apply to any given collection of community types to determine if they share an assembly archetype or not. For example, this synthesis operation answers the question we posed above: the community types *H*_*A*_, *H*_*B*_ and *H*_*C*_ in Fig. 1e share the assembly archetype “a species composition coexist if either species 1 or 2 are present”, while *H*_*D*_ does not. We combine our formalism with experimental data of microbial communities and synthetic data generated from simple population dynamics models to provide evidence of conditions leading to the existence of assembly archetypes. We corroborate these results by studying assembly archetypes using clinical data on the human gut microbiota across healthy and unhealthy hosts.

### Synthesizing assembly archetypes

Denote by Σ = {1, 2, …, *S*} the potential species pool and by Θ the set of different environmental contexts (“environments”, from hereafter) under which we can assemble a community. The *community type H*_*θ*_ of the given species pool Σ under the environment *θ* ∈ Θ is the collection of all species compositions that can coexist under that environment (Methods). We can conveniently visualize a community type as a hypergraph [25] that, in our analogy to limb types, should be regarded as its “skeleton” (Fig. 2a). To describe the assembly rule *M*_*θ*_ of the community type *H*_*θ*_ it is necessary to choose a language and a syntax. Community types are discrete objects. Therefore, boolean functions *M*_*θ*_ : {0, 1}^*S*^ → {0, 1} provide a compact and convenient language to describe them. An assembly rule *M*_*θ*_(*z*) takes as input a binary vector *z* ∈ {0, 1}^*S*^ representing a potential species composition, where its *i*-th entry satisfies *z*_*i*_ = 1 if the *i*-th species is assembled, and *z*_*i*_ = 0 otherwise. For example, *z* = (1, 0, 1) represents the species composition {1, 3}. The assembly rule outputs *M*_*θ*_(*z*) = 1 if the species composition *z* can coexist in the environment *θ*, and *M*_*θ*_(*z*) = 0 otherwise. Therefore, the assembly rule *M*_*θ*_ acts like a filter characterizing the conditions that a species composition needs to satisfy to coexist in environment *θ* [14, 26].

**Figure 2:**
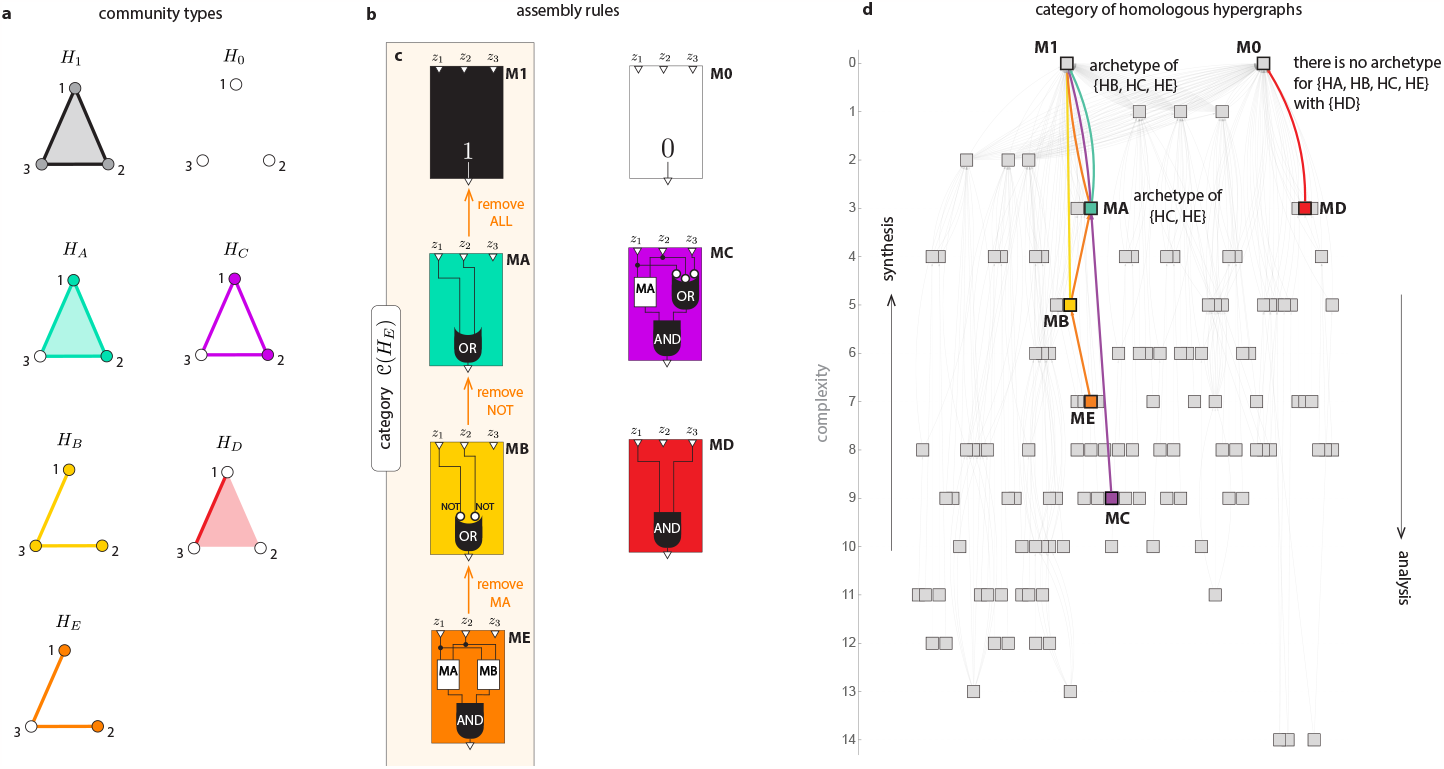
The category of homologous hypergraphs allows synthesizing assembly archetypes in ecological communities. The mathematics of Category Theory allows defining an abstract “synthesis operation” to identify if a collection of community types share an assembly archetype or not. **a**. The community types observed in the environmental context *θ* ∈ Θ can be conveniently visualized as coexistence hypergraphs *H*_*θ*_. In a coexistence hypergraph *H*_*θ*_, vertices correspond to species (circles) and hyperedges to species composition that can coexist in environment *θ*. Filled (resp. empty) circles represent species that survive (resp. cannot survive) alone in the environment. Filled areas represent hyperedges (i.e., species compositions that coexist). For example, the hypergraph *H*_*D*_ = {{1, 3}, {1, 2, 3}} is represented by a line and the interior of a triangle. Coexistence hypergraphs with different color represent different environmental contexts (colors as in Fig. 1e). In particular, *H*_1_ is the complete hypergraph (i.e., all species compositions can coexist), and *H*_0_ is the empty hypergraph (i.e., no species composition can coexist). **b**. Each hypergraph *H*_*θ*_ has an associated *assembly rule M*_*θ*_, describing the conditions that a species composition needs to satisfy to coexist in environmental context *θ*. In this work, we use logic circuits written in the Conjunctive Normal Form as the language and syntax to describe assembly rules. An assembly rule takes as input a binary vector *z* = (*z*_1_, …, *z*_*S*_) ∈ {0, 1}^*S*^ representing a potential species composition, where *z*_*i*_ = 1 only if species *i* is assembled. Then, the assembly rule outputs *M*_*θ*_(*z*) = 1 only if the species composition *z* coexists in the environmental context *θ*. **c**. The category 𝒞(*H*_*E*_) of homologous hypergraphs for the community type *H*_*E*_ contains as objects its assembly archetypes (squares) and their organization as morphisms (arrows). These morphisms correspond to removing variables, logic gates, or whole clauses from one archetype to obtain a simpler one. In this example, its category has four objects {*M*_*E*_, *M*_*B*_, *M*_*A*_, *M*_1_} and three (prime) morphisms. **d**. The category of homologous hypergraphs for all 128 community types *H*_*θ*_ of *S* = 3 species, with objects visualized as squares and morphisms as lines. The resulting categories form a hierarchy. This hierarchy allows defining two operations: synthesis (moving up in this hierarchy) and analysis (moving down in this hierarchy). An assembly archetype *M*\* exists for a collection {*H*_*θ*_, *θ* ∈ Θ} of community types if we can synthesize each community type into the same assembly rule by moving up in the hierarchy. In the panel, *M*_*A*_ is the assembly archetype of {*H*_*C*_, *H*_*E*_}, and *M*_1_ is the assembly archetype of {*H*_*A*_, *H*_*B*_, *H*_*E*_}. Some collections of community types, such as {*H*_*A*_, *H*_*B*_, *H*_*C*_, *H*_*D*_}, do not have an assembly archetype.

To choose a syntax for writing assembly rules, we aim to mimic how more complex assembly archetypes of limb types are constructed by adding constraints to an existing archetype. For example, in Fig. 1a-b, we obtained the assembly archetype “tetrapod with digits” starting from “tetrapod” and adding the constraint “distal bones are digits”. We choose logic circuits written in the minimal Conjunctive Normal Form syntax to obtain a similar behavior for assembly rules (Methods). An assembly rule thus reads as a set of clauses joined by an AND (∧) gate (Fig. 2b). Each of those clauses consists of a combination of the variables {*z*_1_, …, *z*_*S*_} with OR (∨) and NOT (¬) gates. To coexist, a species composition must satisfy each and every clause. Note that adding variables, logic gates, or complete clauses to an existing assembly rule will generate a more complex assembly rule, while removing them will simplify it. This fact suggests defining the *complexity κ*(*M*_*θ*_) of rule *M*_*θ*_ as the minimum number of variables and logic gates needed to write it. In our example of Fig. 1, the community type *H*_*B*_ corresponding to the environment *B* has *M*_*B*_(*z*) = (¬*z*_1_ ∨ ¬*z*_2_) as assembly rule (yellow in Fig. 2b). This assembly rule has only one clause and it shows that, in the *B* environment, a species composition coexists if and only if “either species 1 or species 2 are not assembled”. The complexity of this assembly rule is *κ*(*M*_*B*_) = 5 because it uses two variables and three logic gates. Finally, for each community type *H*_*θ*_, we build a category 𝒞(*H*_*θ*_) encoding its assembly archetypes and how they are organized (Methods). Specifically, the objects in 𝒞(*H*_*θ*_) are assembly rules that are archetypical in the sense that they are the Pareto-optimal descriptions of the community type *H*_*θ*_ in terms of accuracy and complexity. Morphisms in 𝒞(*H*_*θ*_) consist in removing logic gates or entire clauses from an assembly archetype to obtain another simpler one, providing the structure that organizes them. In general, different community types give rise to categories having different objects and morphisms.

To illustrate how our approach works, consider again the community type *H*_*E*_ for a pool of *S* = 3 species shown in Fig. 2a. Figure 2c depicts its corresponding category 𝒞(*H*_*E*_). This category has four objects {*M*_*E*_, *M*_*B*_, *M*_*A*_, *M*_1_}, representing the assembly archetypes of *H*_*E*_. The (prime) morphisms in 𝒞(*H*_*E*_) are 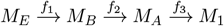, indicating how its assembly archetypes are organized (orange arrows in Fig. 2c). The first assembly archetype *M*_*E*_(*z*) = (¬*z*_1_ ∨ ¬*z*_2_) ∧ (*z*_1_ ∨ *z*_2_) of complexity *κ*(*M*_*E*_) = 7 has two clauses and describes *H*_*E*_ exactly (i.e., each species composition *h* ∈ *H*_*E*_ satisfies *M*_*E*_(*z*(*h*)) = 1, and vice versa). The first prime morphism *f*_1_ removes the clause (*z*_1_ ∨ *z*_2_) from *M*_*E*_ to obtain the second assembly archetype *M*_*B*_(*z*) = (¬*z*_1_ ∨ ¬*z*_2_) of complexity *κ*(*M*_*B*_) = 5. The fact that *M*_*B*_ is the second archetype means that there is no assembly rule that is more accurate for describing *H*_*E*_, except for the first archetype *M*_*E*_ itself. At this complexity level, *H*_*E*_ is best described by the rule “a species composition coexist if either species 1 or 2 is not present”. The second morphism *f*_2_ removes the not gate ¬ from *M*_*B*_ to obtain the third assembly archetype *M*_*A*_(*z*) = (*z*_1_ ∨ *z*_2_) of complexity *κ*(*M*_*A*_) = 3. This third archetype shows that *H*_*E*_ is best described at this complexity level by the rule “a species composition coexist if either species 1 or 2 is present”. The final morphism *f*_3_ removes the clause (*z*_1_ ∨ *z*_2_) from *M*_*A*_ to get a simplest assembly archetype *M*_1_(*z*) = 1 with complexity *κ*(*M*_1_) = 0. This final archetype shows that the best description of *H*_*E*_ at this complexity is simply the rule “all species compositions can coexist”. Note that it is also possible to get the third archetype *M*_*A*_ directly from the first one *M*_*E*_ by composing the two prime morphisms *f*_2_ ° *f*_1_(i.e., by removing the clauses *f*_1_ and *f*_2_ from *M*_*E*_). It is the structure of a category that ensures we can always compose morphisms in this way. The more morphisms we apply to *M*_*E*_, the simpler the assembly archetype we get.

The above categories provide a “synthesis operation” that identifies if an assembly archetype exists for a collection of community types (calculated as the co-product of the union of their categories, Methods). To explain how this works, in Figure 2d, we show the categories 𝒞(*H*_*θ*_) for all the 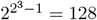 possible community types *H*_*θ*_ of a pool of *S* = 3 species, with objects visualized as squares and morphisms visualized as lines. The resulting categories generate a hierarchy among community types, analogous to that of Fig. 1b for limb types. This hierarchy allows defining two operations: *synthesis* corresponding to moving up in this hierarchy, and *analysis* corresponding to moving down in this hierarchy. For example, the synthesis for the community type *H*_*E*_ consists of starting on the orange square *M*_*E*_ and moving up in the hierarchy *M*_*E*_ → *M*_*B*_ → *M*_*A*_ → *M*_1_ using the morphisms of its category 𝒞(*H*_*E*_) (orange lines in Fig. 2d). We can apply the same process to obtain a synthesis of the other community types *H*_*A*_(green), *H*_*B*_(yellow), *H*_*C*_(purple), and *H*_*D*_(red) using the hierarchy of Fig. 2d, moving using only morphisms of their respective color. Then, an assembly archetype *M*\* exists for a collection {*H*_*θ*_, *θ* ∈ Θ} of community types if we can synthesize each community type into the same assembly rule by moving up in the hierarchy. In other words, an assembly archetype exists if, at some complexity level, the best way to describe the assembly rule of all community types is identical. For example, the community types {*H*_*C*_, *H*_*E*_} have *M*_*A*_(*z*) = (*z*_1_ ∨ *z*_2_) as their assembly archetype. Namely, applying one orange morphism to *M*_*E*_ and one purple morphism to *M*_*C*_ get us to the same assembly rule *M*_*A*_. The assembly archetype for {*H*_*B*_, *H*_*C*_, *H*_*E*_} is *M*_1_(*z*) = 1. Importantly, assembly archetypes not always exist, such as for the community types {*H*_*A*_, *H*_*B*_, *H*_*C*_, *H*_*E*_, *H*_*D*_}. This happens because the first three community types share the assembly archetype *M*_1_(*z*) = 1, but the last community type *H*_*D*_ has the assembly archetype *M*_0_(*z*) = 0 (red in Fig. 2d). That a collection of community types has no assembly archetype means that they do not share a common design principle.

Note we can give a unique label 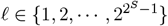 to each of the 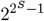 possible assembly rules (or community types) for a pool of *S* species. Then, each environment *θ* ∈ Θ can be associated with the label 𝓁 = 𝓁(*θ*) of the assembly rule *M*_*θ*_ of that environment. Environments with the same assembly rule will have the same label. With a slight abuse of notation, in what follows, we write *M*_𝓁_ and *H*_𝓁_ for the 𝓁-th assembly rule and community type, respectively.

## Results

### Assembly archetypes exist in experimental communities under similar environments

We investigated the existence of assembly archetypes in an experimental microbial community formed by a pool of *S* = 3 bacterial species studied by Friedman et al. [22] (left in Fig. 3). In these experiments, all the possible species compositions were assembled under very similar in vitro conditions. To determine the observed community types, we used the experimental data of species assemblages to probabilistically parameterize a generalized Lotka-Volterra (gLV) model and predict species coexistence (Methods). This method provides an inferred species interaction matrix (left in Fig. 3a), as well as a probability distribution *p*(*θ*) of species’ *effective growth rates θ* = (*θ*_1_, *θ*_2_, *θ*_3_), which phenomenologically represent the total effect of the environment (right in Fig. 3b). Consistent with the experimental design, the inference indicates that the experimental environments are very strongly concentrated into a single value of effective growth rates (black dot in Fig. 3a and Supplementary Fig. 6). Consequently, the community exhibits a unique community type *H*_31_ = {{1}, {2}, {3}} and a unique assembly rule *M*_31_(*z*) = (¬*z*_1_ ∨ ¬*z*_3_) ∧ (¬*z*_1_ ∨ ¬*z*_2_) ∧ (¬*z*_2_ ∨ ¬*z*_3_). This rule itself is the trivial assembly archetype of the community across all experimental similar environments, and it is consistent with the original findings.

**Figure 3:**
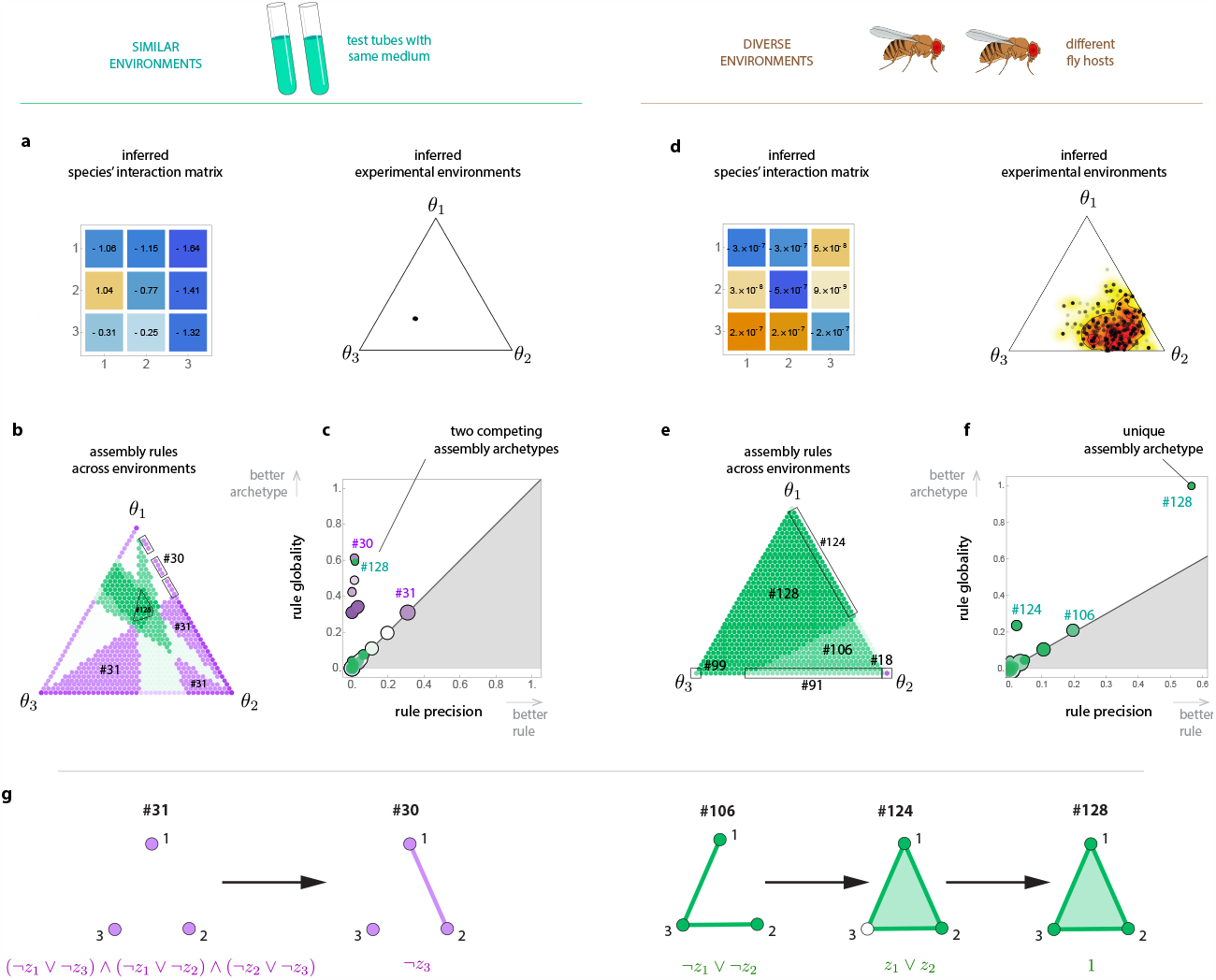
Assembly archetypes in two experimental ecological communities. On the left, a pool of *S* = 3 bacterial species from soil (*Pseudomonas aurantiaca, Pseudomonas veronii* and *Serratia marcescens*) assembled in tightly-controlled environmental contexts [22] (i.e., test tubes with same nutrient medium). On the right, a pool of *S* = 3 species from *Drosophila melanogaster* gut (*Lactobacillus plantarum, Lactobacillus brevis* and *Acetobacter orientalis*) assembled across diverse environmental contexts [27] (different fly hosts). **a**. Interaction matrix and experimental environmental contexts inferred from experimental data. This inference phenomenologically characterizes the environmental context *θ* = (*θ*_1_, *θ*_2_, *θ*_3_) as the species’ effective (intrinsic) growth-rates. Due to the experimental setup, all effective growth rates are positive. For easier visualization, we project them into a ternary plot. The inference indicates that environments are strongly concentrated (black dot). Therefore, across the experimental environmental contexts, the community exhibits a single assembly rule *M*_31_. **b**. Assembling the community across diverse environmental contexts generates not 1 but 24 different assembly rules, each one appearing in a portion of the environment. Here, different colors correspond to different assembly rules. Green represent rules having *M*_128_(*z*) = 1 as archetype, and purple represent rules having *M*_1_(*z*) = 0 as archetype. **c**. The precision *π*(*M*) of rule *M* is the portion of environmental contexts in which it appears. The globality *γ*(*M*) of rule *M* is the portion of environmental contexts in which it is an assembly archetype. The rule with the highest precision is *M*_31_ with *π*(*M*_31_) = 0.31, which coincides with the rule observed in the experimental environment. This most precise rule only occupies less than a third of all environments. Indeed, its globality is also *γ*(*M*_31_) = 0.31. By contrast, there are two assembly archetypes with almost twice the globality: *γ*(*M*_30_) = 0.61 and *γ*(*M*_128_) = 0.59. However, and most importantly, note that there is no global assembly archetype across all environmental contexts. **d and e**. Similar to a and c, but for the community of Drosophila gut microbiota. The inferred environmental contexts (panel d) are more diverse than panel c. These experimental environmental contexts generate six assembly rules (72, 91, 106, 124, 125, 128), and the inferred model predicts that six additional assembly rules are possible (panel e). **f**. Rule *M*_128_ = 1 has both the higher precision and higher globality (*π* = 0.56, *γ* = 0.99), indicating this community has a single and very accurate assembly archetype. **g**. Some of the rules observed in panels a to f and their morphisms.

However, our inferred model suggests that the existence of the above assembly archetype is rather accidental, occurring because of the low diversity of experimental environments. Should the environments be more diverse, the same community would exhibit many different assembly rules. To demonstrate this point, consider the set 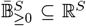 of environments where species have positive effective growth rates and their sum equals those of the experimental environments (i.e., the “total” intrinsic growth rate is kept constant, Methods). If we were to assemble this species pool across 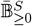, the inference indicates that we would observe not 1 but 24 different assembly rules (Fig. 3b). Each of these assembly rules occupies a fraction of the possible environments. To quantify this fraction, we can define the *precision π*(*M*) of rule *M* as the fraction of all environments in which the rule occurs. More precise rules are those that exactly describe the community types across wider environments. Note that an assembly rule might not exactly describe a community type under some environment, but it might be its assembly archetype. Indeed, each observed assembly rule occupies a portion of the hierarchy of homologous rules, allowing us to determine their archetypes (SI Fig. S12a). Thus, we define the *globality γ*(*M*) of rule *M* as the fraction of all environments under which the rule is an archetype. By definition, *γ*(*M*) ≥ *π*(*M*). A rule has low precision but high globality when it does not exactly describe the assembly of the community across environments but it is an assembly archetype.

When assembling the community across 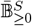, the most precise rule is still *M*_31_, but it has low precision (i.e., *π*(*M*_31_) = 0.31, Fig. 3c). Furthermore, *γ*(*M*_31_) = *π*(*M*_31_), meaning that the validity of *M*_31_ is limited to those environments where it describes the assembly exactly. This result demonstrates that communities can fail to have an accurate assembly rule across environments even in “simple” scenarios (e.g., gLV population dynamics and the environment only changes the species’ intrinsic growth rates).

Despite that no single rule can accurately describe the assembly of this community across environments, we still find it has two accurate assembly archetypes: *M*_30_ = ¬*z*_3_ and *M*_128_ = 1 (Fig. 3c). The first assembly archetype *M*_30_ is twice as global than rule *M*_31_, it is simpler (*κ*(*M*_30_) = 2), and it is easier to interpret (i.e., a composition coexist as long as species 3 is not present). Actually, *M*_30_ is an archetype for rule *M*_31_(purple in Fig. 3g and SI Fig. S12a). The second archetype *M*_128_ has very similar globality to *M*_30_(Fig. 3c). The assembly archetype *M*_128_ occurs only for environments where the effective growth rate of species 1 is the largest among all (green in Fig. 3c). Otherwise, the community has *M*_30_ as the assembly archetype (purple in Fig. 3c). This result is important because it shows that, even in simple scenarios, communities might not exhibit a unique design principle or assembly archetype when assembled across diverse environments. Therefore, given the expected high diversity of environments occurring in nature, this last result may suggest that finding assembly archetypes is unlikely under natural settings.

### Assembly archetypes exist in experimental communities under diverse environments

To investigate if assembly archetypes can also exist in experimental communities assembled across diverse environments, we revisit a community of commensal bacteria in *Drosophila melanogaster* gut studied by Gould et al. [27]. This study experimentally assembled different species compositions fed into different fly hosts (each host representing one particular environment). Using these in vivo experimental data, we inferred again a species interaction matrix (left in Fig. 3d) and a probability density function *p*(*θ*) of effective growth rates (right in Fig. 3d). These inferred effective growth rates are more diverse than those for the in vitro community. Consequently, the in vivo community adopts 6 different assembly rules {*M*_72_, *M*_91_, *M*_106_, *M*_124_, *M*_125_, *M*_128_}, with rule 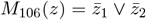 being the most frequent one. Then, as before, we used the inferred model to study the assembly rules that this community can adopt across the even more diverse environments given by 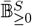 (Fig. 3e). We found that the community adopts 6 additional assembly rules. The observed assembly rules can be located in the hierarchy of homologous assembly rules, allowing us to synthesize their assembly archetype (SI Fig. 12b). Across these diverse environments, the rule *M*_128_(*z*) = 1 is a single remarkably global assembly archetype (*γ*(*M*_128_) = 0.99, Fig. 3f). That is, *M*_128_ is an almost perfect assembly archetype for this species pool, even when assembled across diverse environments. Indeed, note that *M*_128_ is an archetype for rule *M*_106_, which is the most frequent assembly rule observed in the experiments (green in Fig. 3g and SI Fig. S12b). This result provides evidence of the existence of assembly archetypes in experimental communities assembled across diverse environments. The question is why the in vivo community can have an assembly archetype across diverse environments, whereas the in vitro community does not.

### Communities with structured interactions render assembly archetypes

To address the above question, we studied a simple niche model of *S* = 3 competing species (Methods). The niche we consider is a circle (Fig. 4a). Each species occupies a location in this niche, with species located closer competing more strongly. Under these assumptions, the question becomes: are there configurations of species locations that render assembly archetypes more likely when assembling the community across diverse environments? Because the niche is invariant to rotations, we can fix the location of one species without loss of generality. We can then let the other two species’ locations vary across the niche. For each of those locations, we calculate the assembly rules generated when assembling the community across different environments represented by different intrinsic growth rates.

**Figure 4:**
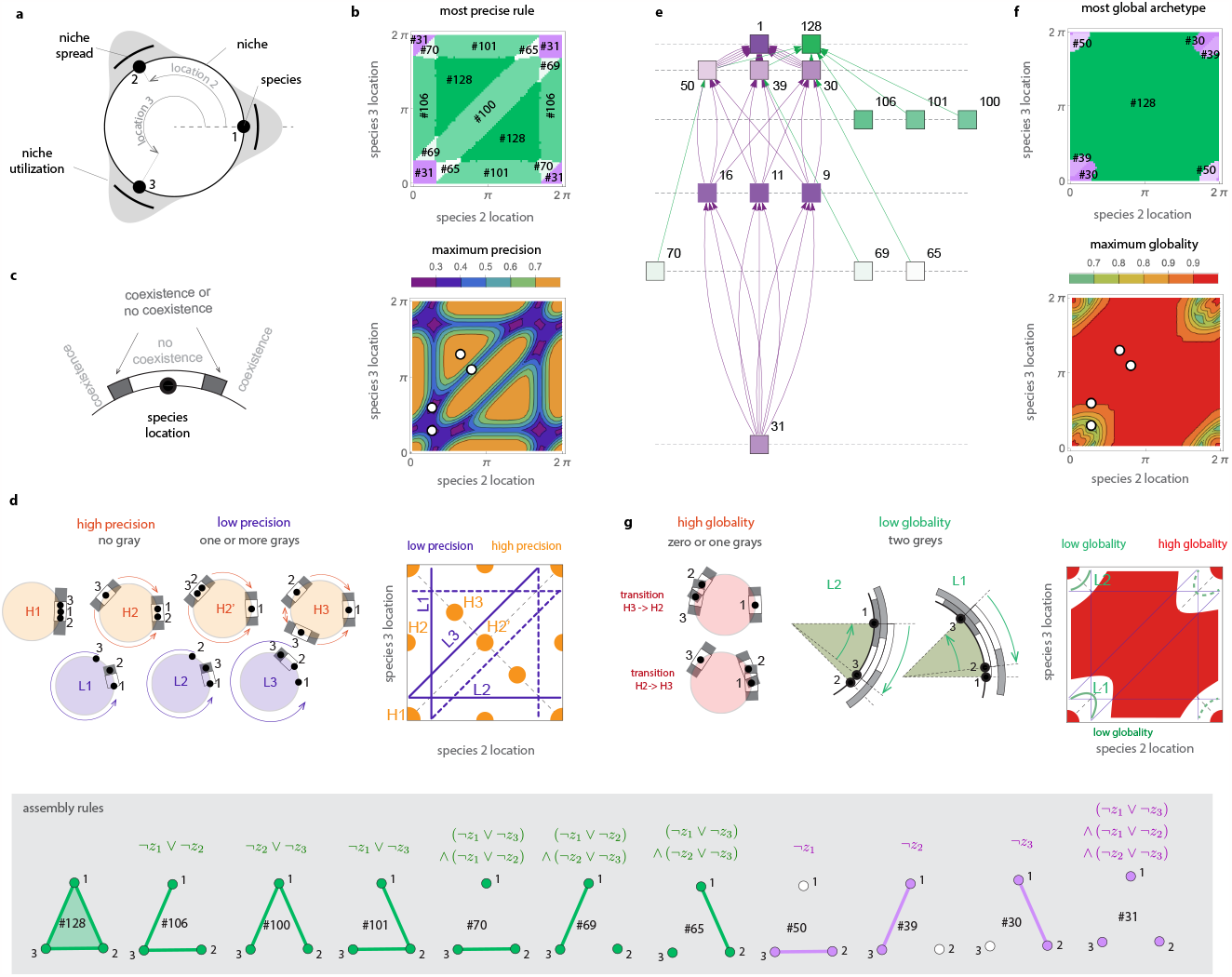
Structured species interactions render assembly archetypes more likely. Results are for a niche model and a pool of *S* = 3 competing species assembled across different environments represented by different intrinsic growth rates chosen uniformly at random from 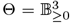. The gray panel at the bottom of the figure shows community types (bottom) with their corresponding assembly rule (top) and their associated label. **a**. Sketch of the niche model, with species located closer competing more strongly. Without loss of generality, we fix the location of the first species at the origin. **b**. Label of the most precise rule (top) and its precision *π*_max_ = max_*θ*_ *π*(*M*_*θ*_) (bottom) across all configurations of species locations. **c**. Competition creates three zones around each species location. The intermediate zone is shown in gray. **d**. Up to symmetry, four configurations H1 to H3 of species locations have high precision (orange). Up to symmetry, configurations with low precision arrange in three lines L1 to L3 (blue). Dashed lines indicate the two symmetry axes. The four configurations with high precision arrange species into one (H1), two (H2 and H2’), and three groups. No species is in an intermediate (grey) zone. Here, species (strongly) compete only with others in the same group. Species can move in the niche direction of the arrows as long they do not enter into an intermediate zone, creating the orange blobs in panel b. For example, in H2, species 3 can approach the group {1, 2}; in H2’, the group {2, 3} move approaching species 1. **e**. Category of homologous hypergraphs observed across all configurations. **f**. Label of the most global assembly archetype (top) and its globality *γ*_max_ = max_*θ*_ *γ*(*M*_*θ*_) (bottom) across all configurations of species locations. **g**. Assembly archetypes are very accurate even if one species is in an intermediate zone (left), thus covering most of the possible configurations of species locations (red in the right panel). Archetypes with low globality occur when two species are in the same intermediate zone. Up two symmetries, there are two such configurations (L1 and L2 in the middle panel). Even in that case, the globality of archetypes is much higher than the precision of assembly rules.

Figure 4b shows the most precise rule found at each species configuration (top) and the maximum precision *π*_max_ that such rule attains (bottom). Configurations where rules attain high precision occur as (orange) blobs and configurations with low precision occur as (blue) lines. Note also the two axes of symmetry in this panel. These blobs and lines can be explained by observing that competition between species creates three zones with different coexistence outcomes around each species’ location (Fig. 4c). First, close to the species location, a zone of no coexistence because strong competition causes competitive exclusion. Second, far from the species location, a zone of coexistence is produced by weak competition. Finally, an intermediate zone between the first two zones where coexistence or no coexistence outcomes are possible depending on the environment where species are assembled. The number of species located in the intermediate zones controls the precision of assembly rules, as shown below.

Assembly rules with high precision occur only when no species is located in an intermediate zone (i.e., when coexistence outcomes are rather environment independent). Up to symmetry, there are four different configurations producing assembly rules with high precision (orange in Fig. 4d). These configurations structure species into one (H1), two (H2 and H2’), or three (H3) groups, where species strongly interact only with other species in their same group. Up to symmetry, there are three lines characterizing configurations with low precision (blue in Fig. 4d). These configurations have in common that at least one species is in an intermediate zone. For example, line L1 characterizes the configurations where species 2 is located in the intermediate zone of species 1, while species 3 location can change across the niche. Configurations at the intersection of two lines occur when two species are located in an intermediate zone, producing assembly rules with very low precision. Importantly, note that the precision of an assembly rule rapidly deteriorates when species are not exactly structured into groups because one species enters an intermediate zone. For example, assembly rules attain a high precision *π*_max_ ≥ 0.7 in the configuration H3 of Fig. 4e. However, if species 2 moves away from species 1 to its intermediate zone, the precision drops by half to *π*_max_ ≈ 0.35.

To calculate the assembly archetypes we build the category of homologous hypergraphs observed across the different species configurations (Fig. 4e). Figure 4f shows the most global assembly archetype found at each species configuration (top) and the maximum globality *γ*_max_ that it attains (bottom). Because globality is at least as high as precision, rules with high globality —and thus accurate assembly archetypes— are also more likely when species are structured into groups. But importantly, accurate archetypes can exist even when this condition is not exactly satisfied. For example, accurate archetypes exist when one species is located in an intermediate zone, such as in the transition from configuration H3 to configuration H2 of Fig. 4g. The configurations with the low globality occur when two species are located in intermediate zones (green in Fig. 4g). However, even in such cases, an assembly rule’s globality remains much higher than its precision (i.e., *γ*_max_ ≥ 0.61 for all species configurations, compared to some configurations having *π*_max_ = 0.2).

Changing the niche’ “size” using a niche spread parameter changes the species’ competition strength. For example, decreasing the niche spread makes the niche “bigger”. In turn, this decreases competition, makes the intermediate zones smaller, and the blue lines of species configurations with low precision become thinner (SI Fig. 14). In this case, because a big circle looks like a line close to any of its points, the neighborhood of the origin will also behave as if the niche was linear (SI Fig. 14a). By contrast, increasing the niche spread will increase the intermediate zones, making the blue lines thicker, overall decreasing the precision and globality of assembly rules (SI Fig. 14b-c). Increasing the niche spreads also tends to increase the number of different observed assembly rule for a species configuration. In general, the largest difference between the globality and precision of an assembly rule occurs at the intersection of the lines L1 and L3 (bottom row of SI Fig. 14). That is, the biggest advantage of assembly archetypes over assembly rules occurs when one species is located in the intermediate zone of each species. We find similar results in a niche model with predator-prey and mutualistic interactions (SI Fig. 15 to 17).

### Assembly archetypes of the human gut microbiota under health and disease

The gut microbiota is the community of bacteria residing in our gastrointestinal tract. It plays a fundamental role in human health and disease [28]. Here, we leverage highly-resolved clinical data analyzed by Gibson et al. [29] to ask if the human gut microbiota has assembly archetypes when assembled across healthy and unhealthy (specifically, with ulcerative colitis) hosts. Specifically, we are interested in understanding similarities and differences under healthy and unhealthy conditions in terms of assembly rules, not in terms of the identities of the taxa assembled in each condition. Thus, we consider a pool of *S* = 16 taxa found in both conditions (Methods). For this taxa pool in healthy and unhealthy hosts, previous work [29] has shown that its dynamics can be explained by: (1) two different structured interaction matrices (Fig. 5a-b), and (2) two different probability distributions for taxa’s effective growth rates (taxa-average [0.15, 0.84] for health, and [0.28, 0.96] for unhealthy, Fig. 5c).

**Figure 5:**
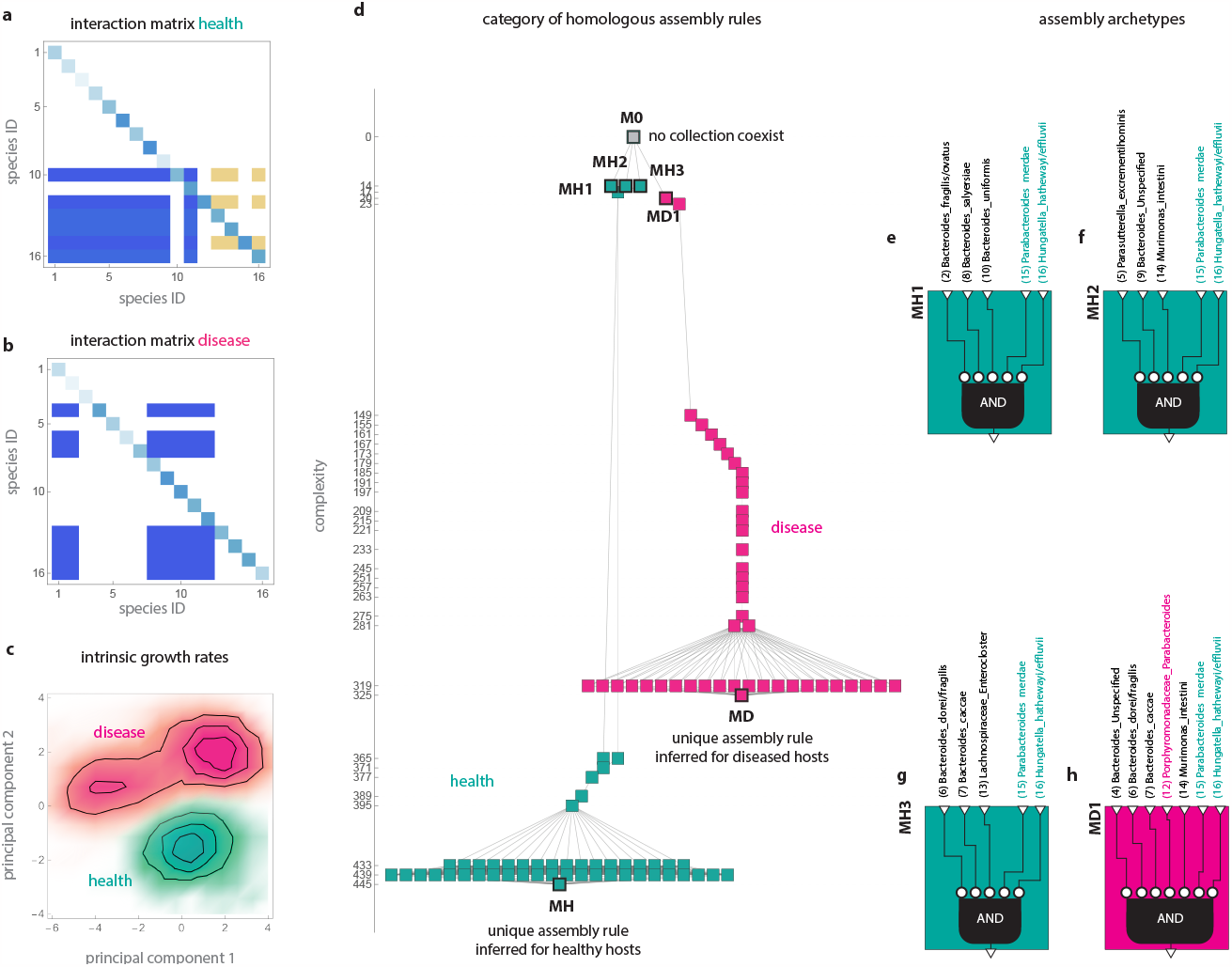
Assembly archetypes in the human gut microbiota under health and disease. Results are for *S* = 16 species shared by the human gut microbiota under health and disease (ulcerative colitis). We use data and inference from [29]. **a-c**. The result of the inference done in Ref.[29] indicates that the gut microbiota dynamics in health and disease across subjects can be explained by two unique, highly structured interaction matrices, one for health (panel a) and one for disease (panel b). Here, blue indicates negative interactions and yellow positive ones. Explaining the microbiota dynamics requires two probability distributions for species’ effective growth rates, one for health (green in panel c) and the other for disease (pink in panel c). This panel shows the two principal components of these two probability distributions for easier visualization. Both distributions show a substantial variance in species’ effective growth rates representing differences in the host’s environmental contexts (average variance across species of 0.026 for health and 0.034 for disease). **d**. Remarkably, the inference indicates that assembling this community across healthy hosts results in a unique assembly rule *M*_*H*_ with complexity *κ*(*M*_*H*_) = 445. Similarly, assembling the community across unhealthy hosts results in a unique assembly rule *M*_*D*_ with complexity *κ*(*M*_*D*_) = 325. We then applied our formalism to investigate if both assembly rules share an assembly archetype, resulting in a category of homologous assembly rules (Methods). We find that the only assembly archetype they share is *M*_0_(*z*) = 0 of zero-complexity, indicating that their assembly is similar only at this lowest complexity level. **e to h**. Other assembly archetypes illuminate the similarities and differences in the assembly of the human gut microbiota under health and disease. The healthy gut has three assembly archetypes of low complexity (MH1, MH2, and MH3) of complexity 14. The unhealthy gut has one assembly archetype of complexity 20. In both health and disease, their archetypes indicate that, to coexist, a species composition must not contain taxa 15 (*Parabacteroides merdae*) and 16 (*Hungatella hathewayi/effluvii*). In the healthy gut, the coexistence of species composition requires that either species {2, 8, 10} (MH1) or (5, 9, 14} (MH2) or {6, 7, 13} (MH3) are not present. Under disease, coexistence becomes possible under stronger conditions, in particular requiring that taxa 12 (*Porphyromonadaceae Parabacteroides*, pink) is not present.

We found that assembling this community across healthy hosts will generate a single community type *M*_*H*_, and thus a single assembly rule *M*_*H*_ of complexity *κ*(*M*_*H*_) = 445. This result is consistent with previous findings indicating that the human gut has remarkably similar dynamics across healthy hosts [30]. Furthermore, the inference indicates that the community also adopts a single (but different) community type *H*_*D*_ when assembled across unhealthy hosts with a rule *M*_*D*_ with complexity *κ*(*M*_*d*_) = 325. That the community adopts a single community type when assembled across healthy or unhealthy hosts is consistent with our previous findings indicating that structured species interactions render more likely the existence of assembly archetypes.

To understand the similarities between the healthy and unhealthy assembly rules, we built the category of homologous assembly rules for them (Methods and Fig. 5d). This category shows that similarities are very scarce: they only share the archetype *M*_0_(*z*) = 0 of zero complexity. The only similarity between health and disease is that, when assembled, most taxa compositions tend not to coexist.

Above the zero-complexity level, the healthy and unhealthy gut assemblies differ. Note these differences are exactly encoded by their assembly rules *M*_*H*_ and *M*_*D*_. But the high complexity of these assembly rules makes their differences challenging to understand. We can circumvent this challenge by comparing their simplest assembly archetypes of non-zero complexity. Our framework synthesizes three healthy assembly archetypes {*M*_*H*1_, *M*_*H*2_, *M*_*H*3_} of complexity *κ* = 14. These three assembly archetypes coincide in that a necessary condition for the coexistence of a taxa composition is that neither *Parabacteroides merdae* (taxa 15) nor *Hungatella hathewayi/effluvii* (taxa 16) is present (green taxa in Fig. 4f-h). In addition, these healthy assembly archetypes indicate that coexistence in the healthy gut requires that at least one of three taxa trios are absent (black taxa in Fig. 4e-g).

Our framework also synthesizes one single assembly archetype *M*_*D*1_ for unhealthy hosts of complexity *κ* = 20 (pink in Fig. 4d). This assembly archetype indicates that, in disease, coexistence requires that: (1) taxa 15 and taxa 16 are not present (green taxa in Fig. 4h); (2) that a quartet of taxa is not present (black taxa in Fig. 4h); and (3) that *Porphyromonadaceae Parabacteroides* is not present (taxa 12, Fig. 4h). While coexistence in healthy hosts also require Condition 1 above, Condition 2 is stronger than in healthy hosts by combining more taxa. Importantly, Condition 3 is not required in healthy hosts. This last observation suggests that taxa 12 becomes highly competitive under disease, making coexistence more challenging. Indeed, clinical studies have found that bacteria of the genus *Parabacteroides* and family *Porphyromonadaceae* are implicated in the genesis of ulcerative colitis by depleting the intestine’ mucosal barrier [31, 32].

## Discussion

Assembly archetypes synthesize the different assembly rules obtained when assembling the same species pool across different environmental contexts, revealing what they all fundamentally share. We have studied experimental and clinical microbial communities and provided direct evidence of assembly archetypes in ecological communities. However, unlike the assembly archetypes for limb types, it is challenging to discover assembly archetypes by direct inspection using our human senses. Instead, assembly archetypes are revealed only through the “mathematical synthesis” that the language of Category Theory provides. It is in this other sense that, paraphrasing David Hilbert’s perspective on the use of mathematics [33, Part I], assembly archetypes are idealized objects that “complete” our partial view of reality. Assembly archetypes are also a concrete realization of Rene Thom’s program to understand biological organization from a mathematical viewpoint (see, e.g., [34, pp. 40] and [35]). We also note that previous works have extended the idea of finding “archetypes” to other biological entities such as genes, nucleotides, physiological processes, and behavioral patterns [36, 37]. To our best knowledge, our work is the first application of the notion of archetype to community ecology.

Our study suggests that assembly archetypes are more likely under structured species interactions. More precisely, we found that rules with high precision tend to occur only when species interactions are such that they make coexistence outcomes environment-independent. This conclusion makes sense because a community assembly can achieve high precision only if it remains similar across environments. Importantly, assembly archetypes can occur under more general conditions when species are structured into groups, where species interact strongly with species within their same group and interact weakly with species in different groups. However, additional work is necessary to study the validity of this result in more detailed mathematical models. For example, considering more species with more detailed niche models of species interactions like foodwebs [38] or mutualistic networks [39, 40], with different niche utilization functions, or allow species to have different kinds of interactions. Indeed, using clinical data with *S* = 16 structured taxa, we have shown that our theoretical and experimental results can remain valid under such more realistic cases. Indeed, we conjecture that a “concentration phenomenon” similar to an asymptotic equipartition property may occur: increasing the number of species could render the partition of environments with the same assembly rule to converge into a configuration where one or a few assembly rules occupy most of the space, while the remaining assembly rules occupy a vanishing or zero space. The clinical data are an extreme example of this conjecture because, according to our findings, a single assembly rule occupies the entire environment. If this conjecture is true, it implies that the counter-intuitive process that assembly archetypes can become more likely as the number of species increases.

Assembly rules with high precision are those with high structural stability [19, 25, 41]. That is, the assembly rule of the species pool remains unchanged despite significant environmental changes. Having assembly rules with high structural stability is sufficient for assembly archetypes because an assembly rule’s globality is at least as large as its precision. However, as Fig. 3b illustrates, this condition is not necessary. Assembly archetypes can exist despite changes in the assembly rules as long as these changes “align” with a single archetype. In this sense, the concept of assembly archetype generalizes the notion of structural stability. For limb types, allowing their assembly rules to change is essential for innovation in body plans and for organisms’ evolvability [42, 43]. Importantly, not all changes are possible because the biological substratum constrains the possible “directions” in which assembly rules can change. These constraints are responsible for generating archetypes [2]. We expect an analogous process occurs in ecology at the level of community types. Namely, when subject to environmental changes, the population dynamics of ecological communities should constrain but not eliminate changes in their assembly rules. It remains open to understand how ecological communities in nature achieve a tradeoff between maintaining robustness against environmental changes and allowing changes in their assembly rules to generate innovations such as new species compositions.

As for future work, the idea of “mining” rules from data is a well-established paradigm in machine learning [44], for instance to mine associations in ecological systems [45]. Rule-mining could be combined with our formalism to make the synthesis of assembly archetypes more efficient for larger communities. In our work, we choose as syntax for assembly rules the Conjunctive Normal Form, but our Category-based framework to synthesize assembly archetypes can be used for any other syntax of interest.

More broadly, according to the Piercian viewpoint [46], the whole scientific endeavor can be conceptualized using only two processes: analysis and synthesis. Analysis starts from the general (e.g., the “species” archetype) and dissects it to discover its particular properties (e.g., which species types are more important for a system). Synthesis is the inverse process, starting from the particular (i.e., a collection of types) and aims to identify the general (i.e., their archetype). Types are the partial, fragmented observations of reality that we have access to. Archetypes “glue” these fragmented observations into a coherent whole or general principle —the unity behind the multiplicity. With the unprecedented amount of available data and computational resources, the time is ripe for ecological research to revisit and develop tools of mathematical synthesis [47, 48], as they could hold the the key to discover general design principles that underlie the complex context-dependent systems we observe in nature.

## Acknowledgments

MTA acknowledges the financial support provided by CONACyT grant No. A1-S-13909 and PAPIIT 104915. Funding to SS was provided by NSF grant No. DEB-2024349.

## Competing financial interests

The authors declare no competing financial interests.

## Author contributions

MTA and SS designed the study and wrote the manuscript. HF performed the numerical analysis.

## Data accessibility

The code supporting the results will be archived on Github.

## Methods

Here we give a summary of our methods, referring the reader to the Supplementary Notes for details.

### Community types

Consider a pool Σ = {1, 2,⋯, *S*} of *S* species that can be assembled in the set Θ ⊆ ℝ^*p*^, *p* ≥ 1, of environmental contexts. Let 2^Σ^ denote the power set of Σ (i.e., the set of all subsets of Σ), representing all potential species compositions that can coexist. When a *species composition h* ∈ 2^Σ^ is assembled in an environment *θ* ∈ Θ, we assume that its coexistence is a binary outcome: the composition either coexists or not. For concreteness and following previous works [25, 49, 50], we use the classic notion of “permanence” as criterion of coexistence. Thus, coexistence allows for species whose abundance converges to one or multiple interior equilibria or that exhibit limit cycles or strange attractors. We emphasize, however, that our framework can use any other coexistence criteria as long as it is binary (e.g., requiring that species’ abundance remains above a certain predefined level). We then define the *community type H*_*θ*_ as the set *H*_*θ*_ = {*h* ∈ 2^Σ^|*h* coexists in environment *θ*}. A community type *H*_*θ*_ can be visualized as a *coexistence hypergraph* [25], with vertices corresponding to species and hyperedges corresponding to species compositions that can coexist under that environment (Definition 1 in SI Note 1.1).

### Assembly rules and their realization as logic circuits

A potential species composition *h* ∈ 2^Σ^ can be equivalently represented by a binary vector *z* = *z*(*h*) ∈ {0, 1}^*S*^ using the convention *z*_*i*_ = 1 only if the *i*-th species is included in *h*. Therefore, any community type *H*_*θ*_ can be equivalently represented by a boolean function *M*_*θ*_ : {0, 1}^*S*^ → {0, 1} where *M*_*θ*_(*z*) = 1 only if the species composition encoded by *z* coexist in environment *θ. M*_*θ*_ is the *assembly rule* for *H*_*θ*_ in the sense that it specifies the conditions that a species composition *z* needs to satisfy for coexisting in the environment *θ* [14]. In this case, we also say that *M*_*θ*_ realizes *H*_*θ*_(Definition 2 in SI Note 1.2).

As syntax to write assembly rules we choose logic circuits written in the minimal Conjunctive Normal Form (CNF), SI Note 1.3. An assembly rule *M*_*θ*_ is thus written as a logic circuit combining the variables {*z*_1_, *z*_2_,⋯, *z*_*n*_} with the logic gates `AND` “∧”, `OR` “∨, and `NOT` “¬”. Specifically, the CNF form of an assembly rule reads as 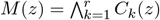, where each *clause C*_*k*_(*z*) is composed only of ¬ and ∨ gates, and *r* ∈ ℕ is the number of clauses. We are interested in the *minimal* CNF representation an assembly rule that uses the minimum number of logic gates and variables (see Example 4 in SI Note 1.3). Very efficient algorithm to compute such minimal CNF representation exist, which are implemented in the standard software packages we use such as Mathematica or PyEDA. A species composition *z* ∈ {0, 1}^*S*^ coexists if and only if it satisfies the clause *C*_*k*_ for all *k* = 1,⋯, *r*.

The convention of writing assembly rules as minimal CNF circuits allows us to define the *complexity κ*(*M*) of the assembly rule *M* as the minimal number of variables and logic gates needed to write it (Definition 3 in SI Note 1.3). The Conjunctive Normal Form imitates our use natural language to describe organisms’ limb types at different complexity levels by adding or removing clauses (SI Fig. S1). More complex assembly rules have more complex clauses or/and a larger number of them.

### The category of homologous community types

We quantify how well a given assembly rule 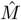 describes a community type *H* using the *prediction error*

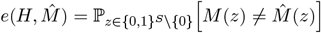

where *M* realizes *H*. Note that 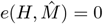 if and only if 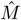 realizes *H*.

Consider now the set ℳ_*S*_ containing all the 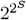 different assembly rules 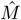 that exist for a pool of *S* species. For each 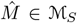, we can calculate its complexity 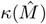 and prediction error *e*(*H, M*). From this data, we identify the assembly rules that are Pareto-optimal in the sense that they provide the best complexity-prediction error tradeoff (Definition 4 in SI Note 2.1). This approach was used before in machine learning for discovering fundamental laws or rules from data, such as Newton’s law of motion [51] or functional responses between species interactions [52]. With this notion, we define the *assembly archetypes* 𝒫(*H*) of *H* as the optimal-Pareto assembly rules that satisfy the additional condition that simpler assembly archetypes must be contained in more complex archetypes, starting with the assembly rule *M* realizing *H* itself as the first archetype (Definition 5 in SI Note 2.1). Each 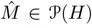 represents the best description of the community assembly rule at the complexity level 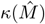. We illustrate this construction in Example 5 of SI Note 2.1 and SI Fig. 2a-d.

From the set 𝒫(*H*) of assembly archetypes for the community type *H*, we build the category 𝒞(*H*) that encodes how the different archetypes can be compared (Definition 6 in SI Note 2.2). Namely, the objects of 𝒞(*H*) are the assembly archetypes of the community type *H*. Morphisms in 𝒞(*H*) represent removing variables, logic gates or full clauses from one archetype to obtain a simpler ones (see SI Fig. 3 for an example of what a morphism represents). We illustrate this construction in Example 6 of SI Note 2.2 and SI Fig. 2e.

### A synthesis operation to calculate assembly archetypes

The constructed categories 𝒞(*H*_*θ*_) for the different community types *H*_*θ*_ allows defining a *synthesis* operation to identify and calculate if an assembly archetype exists for a given collection {*H*_1_,⋯, *H*_*n*_} of community types. To define this synthesis operation, denote the *union category* of all community types as 𝒞(*H*_1_,⋯, *H*_*n*_), see Definition 7 in SI Note 2.3. The objects of this category are the union of the objects of each category 𝒞(*H*_*i*_), and a morphism exists between two objects if it exists in at least one category 𝒞(*H*_*i*_). Then, we define the *assembly archetype M*\*(*H*_1_,⋯, *H*_*n*_) of the community types {*H*_1_,⋯, *H*_*n*_} as the co-product

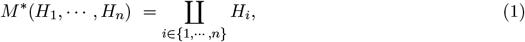

calculated in the category 𝒞(*H*_1_,⋯, *H*_*n*_), if such co-product exists. If the co-product does not exist, then the community types do not have an assembly archetype. See Definition 8 in SI Note 2.4 for details.

### Experimental communities, inference, and estimating community types

We studied three experimental species pools of bacteria based on previously reported experimental data of assemblies of species compositions [22, 27, 29] (see SI Note 3.1 for descriptions). For each species pool, we used the data of different species assemblies to probabilistically parameterize a generalized Lotka-Volterra (gLV) model allowing us to estimate the community types that are likely observed under the experimental environmental contexts. Specifically, for a pool of *S* species, the gLV model takes the form

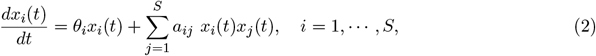

where *x*_*i*_(*t*) denotes the abundance of species *i* at time *t* ≥ 0. The gLV has as parameters the species’ intrinsic growth rates *θ* = (*θ*_1_,⋯, *θ*_*S*_) ∈ ℝ^*S*^ and their interaction matrix *A* = (*a*_*ij*_) ∈ ℝ^*S×S*^. We note that, despite its simplicity, the gLV was found to adequately model the dynamics of two of the three experimental communities we studied [22, 29].

We combined the experimental data of species assemblages with a Bayesian inference method to estimate the gLV parameters under the experimental conditions of each study (SI Note 3.2). For Friedman [22] and Drosophila [27] communities, this resulted in one estimated interaction matrix *Â* and one probability density function *p*(*θ*) for the intrinsic growth rates that phenomenologically represent the diversity of environments in which species are assembled. For gut microbiota [29], we used inferred interaction matrix and probability density function for intrinsic growth rates reported in the original study. In this case, there exists one pair (*Â*_*h*_, *p*_*h*_(*θ*)) for health and another pair (*Â*_*d*_, *p*_*d*_(*θ*)) for disease.

Given a pair (*Â, p*(*θ*)) parametrizing the gLV model, we obtained the observed community types by sampling *θ* ∼ *p*(*θ*) and calculating the coexistence of each species composition using Jansen’s permanence criterion [49, 53] (SI Note 3.3). This resulted in a collection {*H*_*θ*_} of community types that are likely to be observed under the given experimental conditions of each study.

For each experimental system, we construct the set 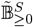 of environments as follows. Let 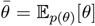 exp denote the expected (i.e., average) value of inferred experimental effective growth rates, where *p*(*θ*) is the distribution inferred from the experimental data. Then we define 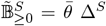, where 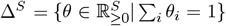 is the -dimensional unit simplex. Therefore, for any vector 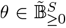, the sum of effective growth rates equals the average effective growth rate observed in the experiment.

### A niche model shows that structured species interactions render assembly archetypes more likely

Our model is an instance of the classic niche model of species competition [54] (see details in SI Note 4). We consider an (abstract) niche that is one-dimensional, finite, and without boundaries, thus topologically equivalent to a circle 𝕊^1^. Each species is assigned a niche *location µ*_*i*_ ∈ 𝕊^1^. We chose a circular niche because it has no distinguished location, so that all species will have an equal number of neighbors on each side. For example, if the niche represents the different resources that can be consumed, a circular niche means there is no special diet. A circular niche also allows us to avoid “edge effects” during the analysis [55]. As noticed before, we can choose the location of the first species as *µ*_1_ = 0 without loss of generality.

To describe the niche use of each species, we use a von Mises distribution centered at the species’ location *µ*_*i*_ and with niche spread 1*/k >* 0. The von Mises distribution is analogous to a Normal distribution on the circle [56]. We choose the niche spread equal for all species. Under the above conditions, the interaction strength of species *j* over species *i* can be calculated as

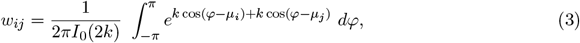

where *I*_0_(*k*) is the modified Bessel function of the first kind of order zero. The value of *α*_*ij*_ depends on the species locations *µ*_*i*_ and *µ*_*j*_. Decreasing the niche spread making *k* → ∞ renders the niche “ bigger” (*k* → 0 renders the niche “smaller”). Note that a sufficiently big circle looks like a line in small neighborhood of any of its points (formally, they are topologically equivalent). Therefore, for small values of the niche spread, the outcomes of our analysis for the circular niche model coincide with the outcomes for a linear niche model.

The dynamics for each species takes the form of the gLV model of (2) with the entry *a*_*ij*_ of the interaction matrix given by *a*_*ij*_ = −*w*_*ij*_. Note this yields *a*_*ii*_ = −1. To fix the species’ intrinsic growth rates *θ* = (*θ*_1_, *θ*_2_, *θ*_3_), we first choose a positive vector 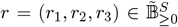 as before, and then set *θ*_*i*_ = *r*_*i*_. In Supplementary Note. 4.3, we describe in details how we extend our niche model to consider predator-prey or mutualistic interactions.

### Synthesizing assembly archetypes in the human gut microbiota

For the pool of *S* = 16 taxa in the human gut microbiota shared under health and disease [29], there exist 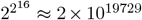 different community types, which is much larger than the estimated number of atoms in the universe. This fact makes it unfeasible to synthesize assembly archetype by constructing the union category 𝒞(*H*_*θ*_) of all possible community types. To circumvent this challenge, we designed a random sampling algorithm that estimates the assembly archetypes for large species pools (SI Note 3.4 and 3.5). In brief, given the calculated community types *H*_*h*_ and *H*_*d*_ that the pool exhibits under healthy and unhealthy hosts, we first calculate their (exact) assembly rules *M*_*h*_ and *M*_*d*_ written in the minimal CNF syntax. Then, our random sampling algorithm estimates their assembly archetypes by randomly removing variables, logic gates, or whole clauses from the exact assembly rules *M*_*h*_ and *M*_*d*_. We describe in detail this algorithm in SI Note 3.3. In SI Fig. 11, we test the performance of this random sampling algorithm showing that it exactly recovers the assembly archetypes in the case of *S* = 3 species.

## Code availability

The code supporting the results will be archived on Zenodo.

## Notes

### Competing Interest Statement

The authors have declared no competing interest.

